# Gut Fungi Possess a Conserved Toxin Immunity Gene of Bacterial Origin

**DOI:** 10.1101/2020.10.15.341461

**Authors:** Syed M. Rizvi, Chengxin Zhang, Peter L. Freddolino, Yang Zhang

**Affiliations:** Department of Ecology and Evolutionary Biology, University of Michigan, Ann Arbor, MI 48109 USA; Department of Computational Medicine and Bioinformatics, University of Michigan, Ann Arbor, MI 48109 USA; Department of Biological Chemistry, University of Michigan, Ann Arbor, MI 48109 USA

**Keywords:** horizontal gene transfer (HGT), bacterial toxin immunity, anaerobic fungi

## Abstract

Prokaryotes and some unicellular eukaryotes routinely overcome evolutionary pressures with the help of horizontally acquired genes. In contrast, it is unusual for multicellular eukaryotes to adapt through horizontal gene transfer (HGT). Recent studies identified several cases of adaptive acquisition in the gut-dwelling multicellular fungal phylum *Neocallimastigomycota*. Here, we add to these cases the acquisition of a putative bacterial toxin immunity gene, PoNi, by an ancient common ancestor of four extant *Neocallimastigomycota* genera through HGT from an extracellular *Ruminococcus* bacterium. The PoNi homologs in these fungal genera share extraordinarily high (>70%) amino acid sequence identity with their bacterial donor xenolog, providing definitive evidence of HGT as opposed to lineage-specific gene retention. Furthermore, PoNi genes are nested on native sections of chromosomal DNA in multiple fungal genomes and are also found in polyadenylated fungal transcriptomes, confirming that these genes are authentic fungal genomic regions rather than sequencing artifacts from bacterial contamination. The HGT event, which is estimated to have occurred at least 66 (±10) million years ago in the gut of a Cretaceous mammal, gave the fungi a putative toxin immunity protein (PoNi) which likely helps them survive toxin-mediated attacks by bacterial competitors in the mammalian gut microbiome.

**Significance:** Adaptation via horizontal gene transfer (HGT) is uncommon in multicellular eukaryotes. Here, we report another *bona fide* case of adaptive evolution involving the horizontal transfer of a bacterial toxin immunity gene from extracellular *Ruminococcus* bacteria to gut-dwelling multicellular fungi. The acquired gene may help the fungi compete against bacterial neighbors in the gut.

## Introduction

Adaptation via horizontal gene transfer (HGT) is common in prokaryotes and some unicellular eukaryotes (Keeling and Palmer 2008), but in multicellular eukaryotes adaptive HGT has historically been considered unusual (Ros and Hurst 2009; Yue et al. 2012; Husnik and McCutcheon 2018). Unlike prokaryotes, eukaryotes do not have the conjugative pili and transformation proteins that facilitate most cases of natural HGT (Ku and Martin 2016). Furthermore, germline sequestration in most multicellular eukaryotes reduces their chances of horizontally acquiring heritable exogenous DNA. As a result, most cases of evolutionarily significant HGT in eukaryotes are prokaryote-to-eukaryote transfers involving unicellular eukaryotes or eukaryotes with intracellular prokaryotic symbionts in their germline cells (Yue et al. 2012; Husnik and McCutcheon 2018), although several notable exceptions have recently been documented. Some of the well-supported exceptions include the HGT-aided colonization of terrestrial environments by the aquatic ancestors of land plants (Yue et al. 2012), the HGT-aided colonization of coffee beans by coffee berry borer beetles (Acuña et al. 2012), and the HGT-aided colonization of mammalian gastrointestinal tracts by anaerobic fungi (Garcia-Vallvé et al. 2000; Murphy et al. 2019).

The *Neocallimastigomycota* phylum, also known as the anaerobic fungi, consists of mycelium-producing multicellular microbes adapted to life in the gastrointestinal tracts of herbivorous mammals (Hanafy et al. 2020). These fungi colonized the mammalian gut during the Cretaceous period, and their subsequent adaptive radiation was aided by multiple horizontal acquisitions of bacterial genes (Murphy et al. 2019; Wang et al. 2019). In addition to abiotic stresses, competition with diverse bacterial species in the gut likely exerted strong evolutionary pressures on these fungi. One such pressure might have been the contact-dependent toxins that are secreted by gut bacteria as a means of microbial warfare to inhibit the growth of other neighboring cells.

Bacteria use cell envelope-associated secretion systems to translocate toxins into neighboring prey cells upon direct physical contact, a phenomenon known as contact-dependent antagonism (Klein et al. 2020). Host-associated bacteria, such as the human pathogens *Vibrio cholerae*, *Salmonella enterica*, *Staphylococcus aureus*, *Bacteroides fragilis*, and *Shigella sonnei*, use contact-dependent antagonism to subvert eukaryotic host cells and/or adjacent prokaryotic competitors in the host microbiome (Ma and Mekalanos 2010; Fu et al. 2013; Cao et al. 2016; Chatzidaki-Livanis et al. 2016; Sana et al. 2016; Anderson et al. 2017). Because the toxins can damage not only prey cells, but also the perpetrator itself, perpetrators possess cognate “immunity proteins” to protect themselves from being damaged by their own toxins (Aoki et al. 2010; Russell et al. 2012).

PoNe is a nuclease toxin domain utilized for interbacterial contact-dependent antagonism by a wide diversity of both Gram-negative and Gram-positive bacteria (Jana et al. 2019). In the genome of the pathogenic bacterium *Vibrio parahaemolyticus*, a PoNe gene was found adjacent to a cognate immunity protein gene (PoNi, previously known as “DUF1911”), and bacterial two-hybrid assays showed that the PoNi protein neutralizes the toxicity of PoNe toxins most likely by directly interacting with them (Jana et al. 2019). In other bacterial genomes, homologs of the PoNi gene were found without an adjacent PoNe toxin gene, suggesting that the PoNi homologs were horizontally acquired and subsequently retained because of their fitness value in conferring immunity against PoNe-mediated antagonism perpetrated by bacterial competitors (Jana et al. 2019).

Although several functional horizontally acquired bacterial genes have been recently identified in anaerobic fungi, such as some antibiotic resistance genes that might protect the fungi against those antibiotics that are indiscriminately released by bacteria (Murphy et al. 2019), to our knowledge no genes that are known to protect against contact-dependent antagonism have been identified. In particular, a recent study reported a substantial rate of HGT between bacteria and anaerobic fungi (Murphy et al. 2019), and even identified in its supplementary materials a protein (GenBank accession: AWI67025.1) homologous to the predicted immunity protein family Imm51 (Zhang et al. 2012), but did not detect any proteins homologous to experimentally verified immunity protein families such as the PoNi family. Furthermore, the Imm51 homolog was found in only one genus (*Caecomyces*) of anaerobic fungi. Due to these two limitations, it is unknown whether bacterial contact-dependent antagonism exerted significant evolutionary pressure on these fungi when they were colonizing and adaptively radiating in the mammalian gut. Here, we identify an extraordinarily conserved and horizontally acquired putative toxin immunity gene, PoNi, in two genomes and multiple transcriptomes of anaerobic fungi. We determined that the fungal PoNi gene, which represents a class of immunity proteins experimentally known to protect against contact-dependent PoNe toxins, was horizontally acquired during the Cretaceous period from an extracellular *Ruminococcus* bacterium. These fungal PoNi proteins share greater than 70% amino acid sequence identity with xenologs in extant bacteria; several additional sequence features add further evidence suggesting that the acquisition of PoNi by anaerobic fungi represents another *bona fide* example of trans-kingdom adaptive horizontal gene transfer to a multicellular eukaryote.

## Results

### HGT of bacterial PoNi gene into fungal genomes

PoNi proteins are bacterial proteins (Jana et al. 2019), and thus the presence of PoNi genes in eukaryotic fungi is a phylogenetic anomaly (indicative of HGT). We identified PoNi genes in the genomes of two anaerobic fungus species, *Anaeromyces robustus* and *Pecoramyces ruminantium*. The PoNi genes are located on scaffold_304:3496-4176 in *A. robustus* (Haitjema et al. 2017) and on c_7180000049905:1398-2081 in *P. ruminantium* (Youssef et al. 2013; Hanafy et al. 2017). Furthermore, blastn search of the NCBI Transcriptomic Shotgun Assembly (TSA) database revealed that mRNA sequences (Supplementary Text S1) of homologs from at least four species of anaerobic fungi are available in the GenBank Nucleotide database: *Anaeromyces contortus* (GGWR01037537.1), *Pecoramyces ruminantium* (GFSU01069946.1), *Caecomyces sp.* Brit4 (GGWS01028851.1 and GGWS01028853.1), and *Piromyces sp.* FS3c (GGXF01056490.1). All of the fungal PoNi homologs share greater than 80% nucleic acid sequence identity with each other (Figure 2B; Supplementary Table S1), as visualized by the sequence alignment in Supplementary Text S1. Similarly, at the amino acid sequence level, the expected translations of these coding sequences all share greater than 77% identity with each other (Supplementary Table S2). We further confirmed that they all contain the DUF1911 domain (which is the old name of PoNi) using NCBI CD-search (Marchler-Bauer et al. 2017), with E-values 3.19E-31, 3.15E-32, 6.74E-33, 6.41E-32, and 1.73E-31 for *A. robustus*, *A. contortus*, *P. ruminantium*, *C. sp.* Brit4, and *P. sp.* FS3c, respectively.

The sequence identity of the fungal PoNi proteins from *P. ruminantium* and *A. robustus* is 84.06% (Table S2). This is much higher than the average global sequence identity (55.74%) for all known *P. ruminantium* proteins (https://www.uniprot.org/uniprot/?query=taxonomy:1987568) and their closest *A. robustus* homologs (https://www.uniprot.org/proteomes/UP000193944). These data suggest that the high sequence conservation of fungal PoNi after HGT is not merely a result of anaerobic fungi evolving relatively recently, but also due to the functional importance of PoNi which mandates a high selective pressure to maintain the conservation of the sequence and function.

### Bacterial donor of fungal PoNi

We next attempted to establish the bacterial donor of the identified fungal PoNi genes. Standard Protein BLAST searches for homologs of the expected translations of fungal PoNi coding sequences revealed that all fungal PoNi homologs are most similar to the same PoNi protein from the same bacterium *Ruminococcus albus* (Supplementary Table S3). The fungal PoNi protein and the *R. albus* PoNi protein share an extraordinarily high amino acid sequence identity given the phylogenetic relationship of the host organisms; the *R. albus* PoNi protein (GenBank protein accession: EGC02125.1) shares an amino acid sequence identity of 73.57%, 72.81%, 73.13%, 75.44%, and 71.37% with the fungal PoNi protein in *A. robustus*, *A. contortus*, *P. ruminantium*, *C. sp.* Brit4, and *P. sp.* FS3c, respectively (Supplementary Table S2). To help visualize the similarity between the fungal and bacterial PoNi proteins, we used C-I-TASSER (Zheng et al. 2019) to create tertiary structure models of PoNi proteins from *A. robustus*, *P. ruminantium*, and *R. albus*. As seen in Figure 1, the two fungal PoNi models (Figure 1A and 1B) are clearly very similar to the *R. albus* bacterial PoNi model (Figure 1C); each shares a TM-score (Xu and Zhang 2010) of 0.97 with the *R. albus* bacterial PoNi model, as calculated by the tertiary structure comparison algorithm TM-align (Zhang and Skolnick 2005).

**Figure 1.**
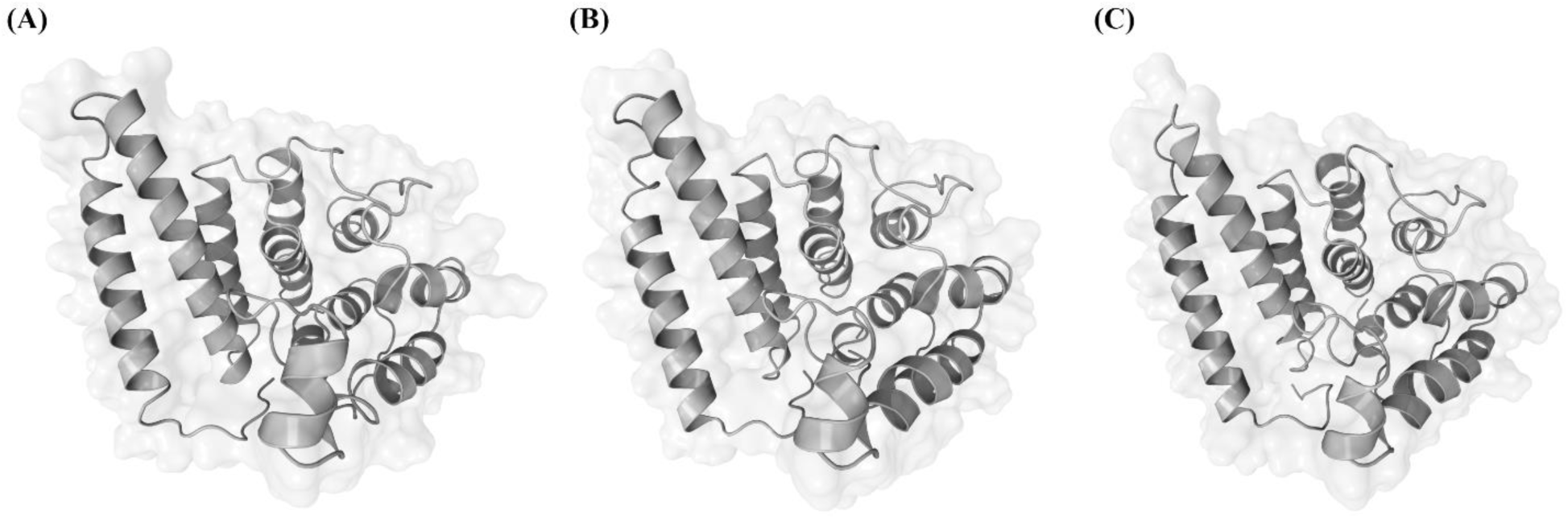
C-I-TASSER tertiary structure models of the PoNi protein from **(A)** *Anaeromyces robustus*, **(B)** *Pecoramyces ruminantium*, and **(C)** *Ruminococcus albus*.

Physical proximity and cohabitation of the same harsh environment may increase the likelihood and utility, respectively, of HGT between microorganisms (Lo et al. 2007; Stewart 2013; Jeong et al. 2019). In our case, anaerobic fungi and *Ruminococcus* bacteria live together in the antagonism-rife mammalian gut (Gordon and Phillips 1998). Furthermore, in addition to sharing the same ecological niche, they might also share the same microniches on the surfaces of food particles within the gut, as suggested by some experimental studies of anaerobic fungi and *Ruminococcus* bacteria (Gordon and Phillips 1998). In particular, *Ruminococcus* bacteria were observed to produce extracellular factors that inhibit the growth of anaerobic fungi on cellulosic substrates, and this was interpreted to be a mechanism that these bacteria evolved to gain a competitive advantage over the fungi. Our current study shows that, in the process of competing for the same microniches as the anaerobic fungi, these *Ruminococcus* bacteria inadvertently increased the fitness of their foes by horizontally providing them a precious toxin immunity gene.

### Fungal PoNi does not arise from bacterial contamination

Contamination from bacteria, which occasionally happens in eukaryotic genome sequencing (Steinegger and Salzberg 2020), is an unlikely explanation for the presence of the bacterial PoNi gene in the fungal genome assemblies. Firstly, the PoNi gene is found in not one but two fungal genomes and in at least four fungal transcriptomes, and the PoNi genes in both of the genomes and all four of the transcriptomes are most closely related to the same species of gut bacterium (*Ruminococcus albus*). Secondly, in each of the two fungal genomes, vertically derived genes are found in the same contig as the PoNi gene. For instance, Figure 2C shows the G+C content around a eukaryotic metallophosphoesterase gene at positions 4000-5500; such humps and valleys in G+C content are signatures of exon-intron architecture in anaerobic fungi (Nicholson et al. 2005). This provides evidence against bacterial contamination because it is unusual for contaminated DNA to be included in the same contig as DNA from the source organism (Richards and Monier 2016; Steinegger and Salzberg 2020). Finally, the presence of mRNA sequences in GenBank provides additional evidence against bacterial contamination, as the employed mRNA sequencing technique only sequenced those RNA molecules that had poly(A) tails (Murphy et al. 2019). Polyadenylation is a routine component of all mRNA processing in eukaryotes, whereas bacteria polyadenylate only a small fraction (less than 3%) of RNA sequences (Régnier and Marujo 2003). Bacterial poly(A) tails also tend to be quite short (less than 20 nucleotides in length), and thus they would not have been efficiently enriched with the usually longer oligo(dT) oligonucleotides used during the sequencing. These differences confirm that the putative fungal PoNi genes are indeed present in genuine regions from the fungi and are not sequencing artifacts from bacterial contamination.

**Figure 2.**
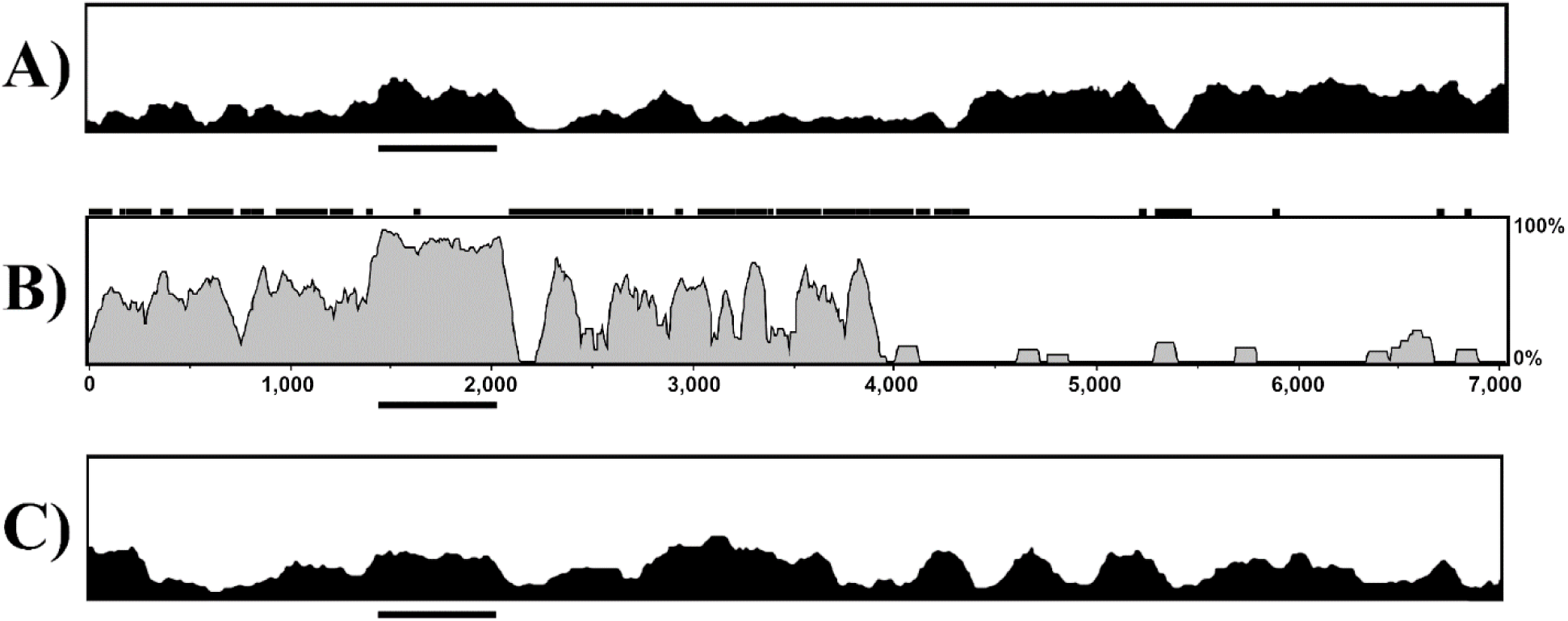
G+C content in a 100 base pair sliding window across the entire 7,034 base pair contig that contains the PoNi gene in *Pecoramyces ruminantium* (**C**) and corresponding region in *Anaeromyces robustus* (**A**). Shared nucleic acid percent identity between the two fungi in a 100 base pair sliding window across this region is shown in the middle (**B**). *X*-axis represents nucleotide position and *y*-axis represents either G+C content (**A** and **C**) or nucleic acid percent identity (**B**). The PoNi gene location is underlined. Small black squares/rectangles above the sequence identity graph indicate the locations of soft-masked repeats in the *Anaeromyces robustus* genome.

### Phylogenetic analysis provides further evidence for HGT of PoNi genes to anaerobic fungi

Phylogenetic incongruence is considered to be the gold standard in HGT detection (Doolittle et al. 2008; Richards and Monier 2016). We constructed a maximum-likelihood phylogenetic tree using the expected translations of the five identified fungal PoNi genes and 292 bacterial PoNi proteins from various bacterial phyla. The tree shows that all five fungal PoNi proteins are nested within *Ruminococcus* bacterial PoNi proteins in the *Firmicutes* bacterial phylum (Figure 3), suggesting that the fungal examples of PoNi genes were horizontally acquired by a single common ancestor, and strengthening the evidence for the HGT event.

**Figure 3.**
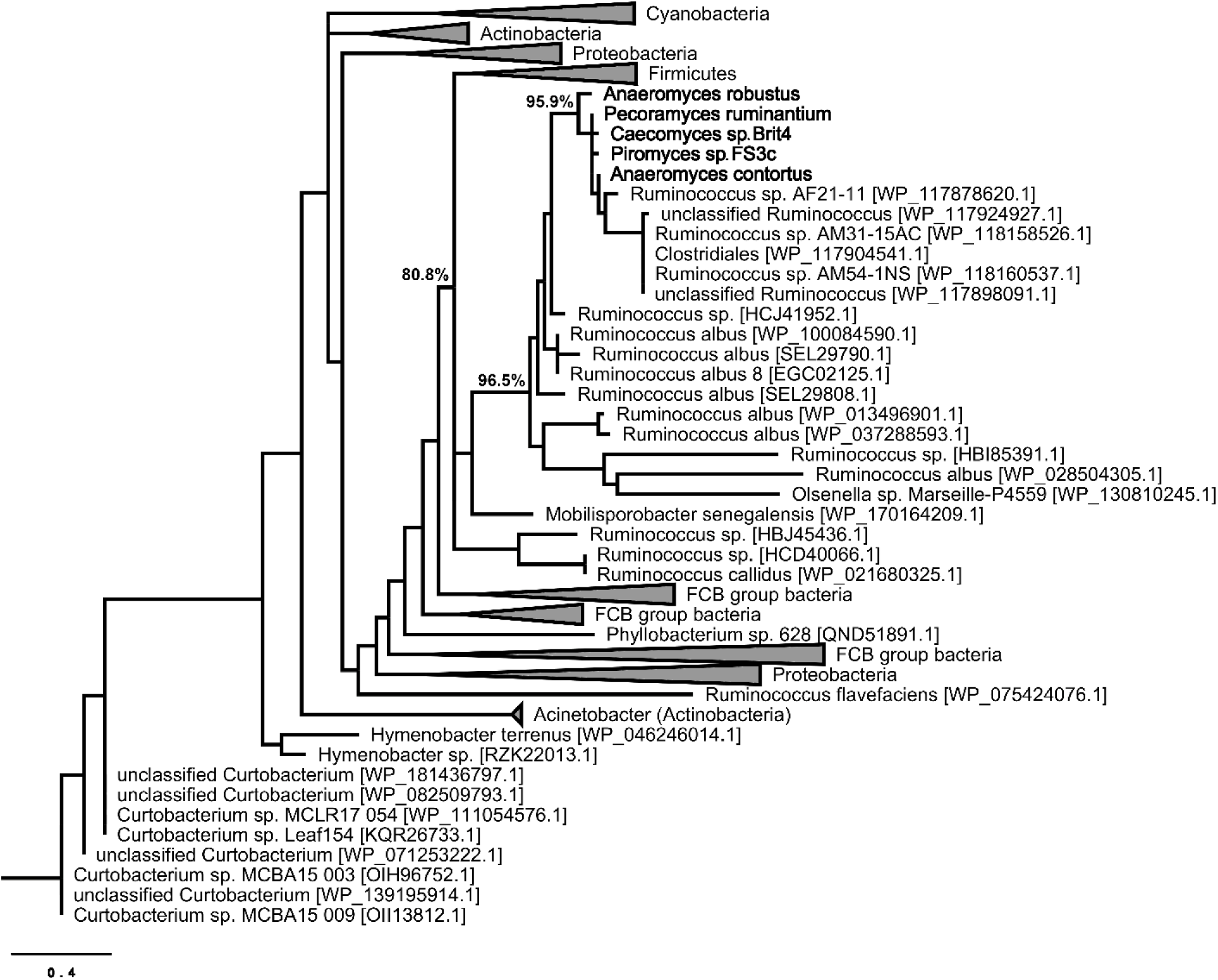
Maximum-likelihood phylogenetic tree of fungal and bacterial PoNi proteins. The fungal proteins are shown in bold. This phylogenetic tree only includes five fungal PoNi proteins, as we could not find PoNi homologs in other fungal lineages. Shimodaira-Hasegawa approximate-likelihood ratio test (aLRT SH-like) values are provided as estimates of branch support for the three most relevant clades next to their nodes (values for all clades are provided in Supplementary Figure S1). GenBank accession numbers are shown in brackets. The tree is drawn to scale, with branch lengths measured in the number of substitutions per site.

### Date of the transfer

The *Anaeromyces*, *Pecoramyces*, *Caecomyces*, and *Piromyces* genera are four cousins that descended from the same ancient gut fungus species (Wang et al. 2019). Given the similarity between the PoNi genes in these genera (Figure 3; Supplementary Tables S1 and S2), we speculate that a common ancestor horizontally acquired a PoNi gene from a *Ruminococcus* bacterium. The bacterium-to-fungus PoNi gene transfer is thus estimated to have occurred at least 66 (±10) million years ago (the Cretaceous period), which is the date for the most recent common ancestor of these four fungal genera (Wang et al. 2019).

## Discussion

Two of the main difficulties associated with validating a prokaryote-to-eukaryote HGT claim are the alternative possibilities of contamination and lineage-specific gene retention (Ku and Martin 2016). Contamination during sequencing has resulted in some high-profile false positive HGT claims in the literature (Richards and Monier 2016), while low (<70%) amino acid sequence identities between xenologs leaves open the alternative explanation of lineage-specific gene retention (Ku and Martin 2016), i.e. that the “horizontally acquired” genes are actually merely the last lone survivors of gene families that were lost in all other eukaryotes (but not yet lost in prokaryotes), creating a phylogenetic distribution pattern that looks superficially like HGT. It should be noted, however, that HGT reports by previous studies (especially those involving older eukaryotic lineages) are not necessarily incorrect due to their lower amino acid sequence identities with bacterial donors. Rather, the low sequence identities fail to preclude the alternative explanation of lineage-specific gene loss, and other arguments (e.g. differences in G+C content) are then required to compensate.

We offer several lines of evidence arguing that we have in fact identified a true inter-kingdom HGT event instead of a case of contamination: the identified PoNi gene is found nested on native sections of chromosomal DNA in multiple eukaryotic genomes, and its RNA has appropriate post-transcriptional modifications for the eukaryotic host genome. Furthermore, the fungal PoNi proteins share extraordinarily high (>70%) amino acid sequence identities with their bacterial donor xenolog, which provides definitive evidence that this was an HGT event rather than lineage-specific gene retention.

Despite the passing of tens of millions of years, the horizontally acquired PoNi gene in anaerobic fungi remains conserved; the high amino acid sequence identity of the fungal PoNi protein with its *Ruminococcus* donor xenolog indicates that natural selection preserved the gene for its functional significance, as pseudogenization is expected in the absence of positive selection (Ros and Hurst 2009; Park and Zhang 2012). Assuming the PoNi gene in these fungi was not co-opted for a novel function, the horizontally acquired toxin immunity gene is predicted to help them survive toxin-mediated attacks by bacterial competitors in the gut microbiome. When the ancestors of anaerobic fungi were colonizing mammalian gastrointestinal tracts in the Cretaceous period, it appears likely that they routinely faced the menace of PoNe-wielding bacterial neighbors, as suggested by the high conservation of the fungal PoNi gene. Horizontal acquisition of the toxin immunity gene served as a quick solution to this menace, allowing the fungi to flourish and adaptively radiate in the antagonism-rife mammalian gut.

Although the eukaryotic recipient in this case was technically a mycelium-producing multicellular fungus, it was nevertheless a relatively simple organism. A previous study on HGT in anaerobic fungi suggested that the weakly protected free zoospore stage in their life cycle provides a possible entry point for horizontal acquisition of heritable exogenous DNA (Murphy et al. 2019).

Our findings revealed how HGT resolved ancient contact-dependent antagonism between newly arrived anaerobic fungal colonists and resident bacteria in the mammalian gut. This antagonism took the form of toxins injected by bacteria into their fungal neighbors, a strategy also known to have been used by ancient bee gut bacteria for interbacterial warfare (Steele et al. 2017). To our knowledge, our current study is the first to implicate PoNe toxins in anti-eukaryotic activity, and our results also highlight the crucial role of HGT in making the anaerobic fungal phylum a successful competitor in the mammalian gut microbiome. The ever-increasing space of available genomic sequences from a broad range of organisms provides an ideal substrate for future identification of other such HGT events.

## Materials and methods

### BLAST

To identify the bacterial donor of the fungal PoNi genes, standard Protein BLAST searches were performed using the blastp (protein-protein BLAST) algorithm with default parameters against the Non-redundant protein sequences (nr) database. The query accession as well as e-value, coverage, and percent identity of the top hits are summarized in Supplementary Table S3.

### Sequence alignment and identity calculation

Pairwise global alignment and sequence identity calculation of nucleic acid sequences (Supplementary Text S1, Supplementary Table S1) and amino acid sequences (Supplementary Text S2, Supplementary Table S2) was performed by NW-align, a simple and robust alignment program for sequence-to-sequence alignments based on the standard Needleman-Wunsch dynamic programming algorithm (Y. Zhang, http://zhanglab.ccmb.med.umich.edu/NW-align). For amino acid pairwise sequence alignments, the BLOSUM62 mutation matrix was used, with gap opening penalty = −11 and gap extension penalty = −1. For nucleic acid pairwise sequence alignments, the default NCBI blastn parameters were used: match = 2, mismatch = −3, gap opening = −5, and gap extension = −2 (Johnson et al. 2008). Sequence identities were calculated based on the length of the longer sequence in each pairwise sequence alignment. The PoNi genomic neighborhood in *A. robustus* (scaffold_304:2098-9132) and *P. ruminantium* (c_7180000049905:1-7034) was aligned by LAGAN (Brudno et al. 2003), and sliding window nucleic acid percent identity and soft-masked repeats visualized (Figure 2B) by VISTA (Mayor et al. 2000). For Supplementary Texts S1 and S2, multiple sequence alignment of PoNi sequences was performed by MUSCLE at the EMBL-EBI website (Edgar 2004; Madeira et al. 2019) and visualized by SeaView (Gouy et al. 2010).

### Tertiary structure prediction by C-I-TASSER

We used C-I-TASSER (Zheng et al. 2019) to create tertiary structure models (Figure 1) of the *A. robustus*, *P. ruminantium*, and *R. albus* PoNi proteins. C-I-TASSER is an extended pipeline of I-TASSER (Yang et al. 2015) and utilizes deep convolutional neural network-based contact maps (Li et al. 2019) to guide the assembly of template fragments into full-length protein tertiary structure by replica-exchange Monte Carlo simulations. The accuracy of the tertiary structure models, measured by estimated TM-score, is 0.95±0.05, 0.94±0.05, and 0.96±0.05 for the *A. robustus*, *P. ruminantium*, and *R. albus* PoNi models respectively. Following strict statistics of structures in the PDB, TM-scores higher than 0.5 assume generally the same fold in SCOP/CATH (Xu and Zhang 2010). Images of the models were created using The Protein Imager (Tomasello et al. 2020).

### G+C content track

G+C content track (Figure 2A and 2C) was obtained from the MycoCosm JGI Genome Browser (Nordberg et al. 2014).

### Phylogenetic tree construction

A maximum-likelihood phylogenetic tree (Figure 3) of five fungal PoNi proteins (Supplementary Text S2) and 292 bacterial PoNi proteins (Supplementary Figure S1) was constructed using PhyML (Guindon et al. 2010). The 292 bacterial PoNi proteins were selected from 12 blastp searches against the Non-redundant protein sequences (nr) database using the *Anaeromyces robustus* PoNi protein as the query (expect threshold = 10^−10^, max target sequences = 100), with each search restricted to a different bacterial phylum (Youssef et al. 2015) in order to get sequences from a diversity of bacterial phyla: *Proteobacteria* (taxid:1224), *Firmicutes* (taxid:1239), *Actinobacteria* (taxid:201174), *Cyanobacteria* (taxid:1117), *Bacteroidetes* (taxid:976), *Spirochaetae* (taxid:203691), *Chloroflexi* (taxid:200795), *Chlorobi* (taxid:1090), *Planctomycetes* (taxid:203682), *Chlamydiae* (taxid:204428), *Deinococcus-Thermus* (taxid:1297), and *Thermotogae* (taxid:200918). No homologs returned for the last five of these phyla. We did not perform HGT index calculation or species tree reconciliation because protein BLAST searches restricted to fungi did not reveal any fungal PoNi homologs other than those in anaerobic fungi. Sequence alignment was performed by MAFFT (Katoh et al. 2018), with gap opening penalty adjusted to 5.0 and the “leave gappy regions” option. Alignment was curated by Gblocks (Castresana 2000), using all options for a less stringent selection (allow smaller final blocks, allow gap positions within the final blocks, and allow less strict flanking positions). PhyML automatically selected the best substitution model (WAG+G) based on BIC criteria. PhyML tested branch support using the Shimodaira-Hasegawa approximate-likelihood ratio test (aLRT SH-like) (Anisimova and Gascuel 2006). The tree was visualized in FigTree v1.4.4 (Rambaut 2018).

## Acknowledgements

This work used the Extreme Science and Engineering Discovery Environment (XSEDE), which is supported by the National Science Foundation grant ACI1548562. This work was supported in part by National Institutes of Health (GM083107, GM136422, AI134678, and OD026825 to Y.Z., and GM128637 to P.L.F.), and the National Science Foundation (IIS1901191 to Y.Z.).

## Author Contributions

Y.Z. conceived the study. C.Z., P.L.F., and S.M.R. designed the experiments. S.M.R. performed the experiments and S.M.R. and C.Z. wrote the initial draft. All authors revised the manuscript and approved the final version.

## Data Availability

The data underlying this article are available in the article and in its online supplementary material.

