## Supplementary texts and tables for "Gut Fungi Possess a Conserved Toxin Immunity Gene of Bacterial Origin"

**Supplementary Material**

**Table of Contents**

**Text S1.** Nucleic acid sequences of the PoNi gene from *A. robustus, A. contortus, P. ruminantium, C. sp.* Brit4*, P.* sp. FS3c, and *R. albus*.

**Text S2.** Amino acid sequences of PoNi proteins from *A. robustus, A. contortus, P. ruminantium, C. sp.* Brit4*, P. sp.* FS3c, and *R. albus*.

**Table S1.** Nucleic acid sequence identities between the PoNi coding sequences, presented as a percent identity matrix.

**Table S2.** Amino acid sequence identities between the PoNi proteins, presented as a percent identity matrix.

**Table S3.** Brief summary of the top hit from BLAST searches, performed using blastp with default parameters against the non-redundant protein sequences (nr).

**Figure S1.** Full, expanded maximum-likelihood phylogenetic tree of fungal and bacterial PoNi proteins. The fungal proteins are shown in bold. This phylogenetic tree only includes five fungal PoNi proteins, as we could not find PoNi homologs in other fungal lineages. Shimodaira-Hasegawa approximate-likelihood ratio test (aLRT SH-like) values are provided as estimates of branch support next to the nodes. GenBank accession numbers are shown in brackets. The tree is drawn to scale, with branch lengths measured in the number of substitutions per site.

### Text S1

**PoNi nucleic acid sequence in *Anaeromyces robustus***

Retrieved from Joint Genome Institute (JGI) MycoCosm website; sequenced by Haitjema et al. 2017.

scaffold_304:3496-4176

ATGAGAGACACTTTAAGAACAAAGGAATATTTTGATACTTTTATTCTTGAAGAATTAGAAGATATTAAAATGTTTGA
AGATAGTATTGAGAATGGGGAAATTGAAGACGAAAGAATTAATTTTATAAAAGACGATATATTAGAAATTAAATTAG
GAATAATTATAGCTAAATATTCCAGAGGAGATCCTATTTGTACAATAAAAAAAGAATTTGAAGATATGATTGATTTT
TTTTGTGAATTTTGGAATGGTGAAATATATGAAGATAATCTTTGGTTTGCTTCTTTGGCCTATCTTTTAAAATTAGA
TGAAGCATTGTTAAAAAGAATAAGAAATAAATTAAAAGAATCTGATACATATGATTATCTTATAGATTTTATACTTA
TAGGTTTTGATGATTCTCAAGATAATTTAAAAATATCATTTCCTCGTCCTTATAAAAAATTATTAAAATGTATTAAT
GGTCAAGATAGGGAAGCATTATTAAAATACTTACGAGGCTGGTATAAAGGTTGTCAAGAAAGCTCATGGTATGATAC
ACACAAAATTAAAGATGATAATTTATATTTCGGATATTGGTGTTTTGAAGCAGGTGCTGTAGCTAAAAGACTTGGTT
TTGAAGACGATGATTTAAAAAATAAACAATATTATCCATATGATATGATTCATTATATTGAATAA

**PoNi nucleic acid sequence in *Anaeromyces contortus***

Retrieved from GenBank (nucleotide accession: GGWR01037537.1)

ATGAAAACGAGAGACACTTTAAGGTCAAAGGAATATTTTGACACTTTTATTCTTGAAGAACTAGAAGATATTAAAAT
GTATGAAGATAGTATTGAGAATGGGGAAATTGAAGAAGATAGAATTGATATAATAAAAGATGATATATTGGAAATCA
AATTAGGAATAATTATAGCTAAATATTCAAGAGGTGATCCTATTAATATAATAAAGAAAGAGTTTGAGGATATGATT
GATTTGTTTTGTGAATATTGGATTGGTGAAATATATGAAGATAATCTTTGTTTTGCTTCTTTAGCATACCTTTTAGC
ATTAGATGATACCTTATTAAATAAAATAAGAAATAAATTAAAAGAATCTGATACATATGATTATCTGATAGATTTTA
TATTGGTAGGTTTTGATGATGATCAAAATGATTCAAAAATATCATTTCCTCGTCCTTATAAAAAATTGTTAAAAAGT
ATAAATGGTAAAGATAGAAATGCTTTTTTGAAGTATTTACGAGGTTGGTATAAAGGTTGTCAGGAAAGCTCATGGTA
CGATACACACAAAATTAAAGATGACAATATATATTTTGGGTATTGGTGCTTTGAAGCAGGTGCTATAGCTAAAAGAC
TTGGTTTTAAAGACGATGATTTAAAAAATGAACAGTATTATCCATACGATATGGTTCATTTTACTGAATGA

**PoNi nucleic acid sequence in *Pecoramyces ruminantium***

Retrieved from Joint Genome Institute (JGI) MycoCosm website; sequenced by Youssef et al. 2013.

c_7180000049905:1398-2081

ATGAAAATGAGAGATACTTTAAGAACAAAGGAATATTTTGACACTTTTATTCAGGAAGAATTGGAAGATATTCATAT
GTTTGAAGATAGTCTTTCGAAAGGAGAAATAGAGGAGGAAAGAATCGATTCGATAAAAGATGAAATATTAGAAATTA
AAATGGGAATAGTTATTGCTAGATATTCAAGAGGGGATTCGATGAATGAACTAAAACAAGAATTTGAAGAAATGATT
GATAGATTTTGTGAATCATGGGACGGTGAAATCTATGAGGATAATCTTTGTTTTGCCTCTTTGGCCTATCTTTTAGG
ATTAGATGATGAGCAATTGAATAGAATAAGAAATAAATTAAAAGAATCGGATACCTATGATTTTCTTATAGATTTTG
TACTTGTGGGTTTTGATGATACTCTGGATGAATCAAAAATATCGTTTCCACGTCCCTATAAACAATTGTTAAAGAGT
ATTAATGGTAAGGATAGAGTTGCTTTCCAGAAATATCTACGAGGTTGGTATAAAGGTTGTCAGGAAAGTTCGTGGTA
TGATACTCATAAAATTGAAGATGATAATCTATATTTTGGATATTGGTGCTTTGAAGCCGGTGCTGTAGCTAAAAGAC
TAGGTTTTAAAGACGATGATTTAAAAAATGAGCAATATTATCCATATGATATGGTTCATTTTACATGA

**PoNi nucleic acid sequence in *Caecomyces sp.* Brit4**

Retrieved from GenBank (nucleotide accession: GGWS01028851.1 and GGWS01028853.1)

ATGAAAATGAGAGACACTTTAAGGACAAAGGAATATTTTGACAAGTTTATTCTTGAAGAACTGGAAGACATTCAAAT
GTTTGAAGATAGTCTTGAGAAAGGGGAAATTGAAAAAGATAGAATCGATTCAATAAAAGACGAAGTGTTACAAATTA
AAGTAGGAATAATTATAGCTAAATATTCAAGAGGTGATCCTATTGATACCATAAAGAAAGAATTTGAAGATATGATT
GATTCGTTTTGTGAATCTTGGGATGGTGAAATATATGAAGATAATCTTTGTTTTGCTTCCTTGGCATATCTTTTAGG
ACTAGATAATACACAGCTAAATAGGATAAGAAATAAATTAAAGGAATCTGATACATACGATTATCTTATAGATTTTG
TACTTGTAGGCGTTGATGATGCTCAGGATAATTCAGAAATATCATTCCCTCGTCCTTATAAACAATTATTAAAATGT
ATTAATGGTGAAGACAAATATGCTTTTAAAAAATATTTACGAGGCTGGTATAAAGGTTGTCAGGAAAGTTCATGGTA
TGATACACACAAAATTGAAGATGATAATTTGTATTTTGGTTATTGGTGTTTCGAAGCAGGAGCTGTAGCCAAAAGAC
TTGGTTTTAAAGATGATGATTTACAAAATGAGCAGTATTATCCATATGATATGGTTCATTTTACTGAATGA

**PoNi nucleic acid sequence in *Piromyces sp.* FS3c**

Retrieved from GenBank (nucleotide accession: GGXF01056490.1)

ATGAGAGATACTTTAAGAACAAAGGAATATTTTGACACTTTTATTCAGGAAGAATTGGAAGATATTCATATGTTTGA
AGATAGTCTTTCGAAAGGAGAAATAGAGGAGGAAAGAATCGATTCGATAAAAGATGAAATATTAGAAATTAAAATGG
GAATAGTTATTGCTAGATACTCAAGAGGGGATTCGATGAATGAACTAAAACGGGAATTTGAAGAAATGATTGATAAA
TTTTGTGATTCATGGGACGGTGAAATCTATGAGGATAATCTTTGTTTTGCCTCTTTGGCCTATCTTTTAGGATTAGA
TGATGAGCAATTGAATAGAATAAGAAATAAATTAAAAGAATCGGATACCTATGATTTTCTTATAGATTTTGTACTTG
TGGGTTTTGATGATACTCTGGATGAATCAAAAATATCGTTTCCACGTCCCTATAAACAATTGTTAAGGAGTATTAAT
GGTAAGGATAGAGTTGCTTTCCAGAAATATCTACGAGGTTGGTATAAAGGTTGTCAGGAAAGCTCGTGGTATGATAC
TCATAAAATTGAAGATGACAATCTATATTTTGGATATTGGTGCTTTGAAGCCGGTGCTGTAGCTAAAAGACTAGGTT
TTAAAGACGATGATTTAAAAAATGAGCATTATTATCCATATGATATGGTTCATTTTGCATGA

**PoNi nucleic acid sequence in *Ruminococcus albus***

Retrieved from GenBank (nucleotide accession: ADKM02000110.1)

ATGCAAATGAGAGATAAATTAAGAAAAAAAGAATACTTCGATACATTCATAGAAGAGGAGATCGAAGACATTCAGAT
GTTTGAAGACAGCCTTGCCGATGGTGAGATCGAAGAAGATAGAATTGATTCAATAAAAGACGAGATACTCCTGATAA
AACTTGGTGTCATTATTGCGAGATATTCAAGATGCGATCCTATAGACGATATCAAGTCGGGATTTGAAGACATGATC
GATATGTTTTGCGAATCATGGGACGGTGGTATATATGAAGATAATCTATGGTTTGCATCATTGGCTTATCTTCTGGG
GCTTGACAGCGCAAAACTTGAAAAAATAAGGAAAAAATTGATGGAAAGCGACACGTATGACTATCTTATTGATTTTA
TCCTATCGGGTACTGAAAGCAAGTTTGACAACAGTAAGATATCTTTCCCGCGTTCATATAAAAAACTGGTCAAAAGC
ATAAATGAGAATGACAAAGAGTCGTTATTAAAATATCTGCGCGGCTGGTATAAAGGCAGTCAGGAAAGTTCCTGGTA
TGATACGCACAAGATCACAGACGACAACCTTTATTACGGCTACTGGTGCTTTGATGCAGGTGCAGTTGCAAAAAGGC
TTGGTCTTGAAGACAGTGACCTGCAAAATGAACAGTACTATCCTTATGATATCGTTCATTTCAGCTGA

**PoNi nucleic acid sequence alignment**

```
                            1
Ruminococcus_albus          ATGCAAATGA GAGATAAATT AAGAAAAAAA GAATACTTCG ATACATTCAT AGAAGAGGAG ATCGAAGACA TTCAGATGTT
Pecoramyces_ruminantium     ATGAAAATGA GAGATACTTT AAGAACAAAG GAATATTTTG ACACTTTTAT TCAGGAAGAA TTGGAAGATA TTCATATGTT
Piromyces_sp._FS3c          ------ATGA GAGATACTTT AAGAACAAAG GAATATTTTG ACACTTTTAT TCAGGAAGAA TTGGAAGATA TTCATATGTT
Caecomyces_sp._Brit4        ATGAAAATGA GAGACACTTT AAGGACAAAG GAATATTTTG ACAAGTTTAT TCTTGAAGAA CTGGAAGACA TTCAAATGTT
Anaeromyces_robustus        ------ATGA GAGACACTTT AAGAACAAAG GAATATTTTG ATACTTTTAT TCTTGAAGAA TTAGAAGATA TTAAAATGTT
Anaeromyces_contortus       ATGAAAACGA GAGACACTTT AAGGTCAAAG GAATATTTTG ACACTTTTAT TCTTGAAGAA CTAGAAGATA TTAAAATGTA

                            81
Ruminococcus_albus          TGAAGACAGC CTTGCCGATG GTGAGATCGA AGAAGATAGA ATTGATTCAA TAAAAGACGA GATACTCCTG ATAAAACTTG
Pecoramyces_ruminantium     TGAAGATAGT CTTTCGAAAG GAGAAATAGA GGAGGAAAGA ATCGATTCGA TAAAAGATGA AATATTAGAA ATTAAAATGG
Piromyces_sp._FS3c          TGAAGATAGT CTTTCGAAAG GAGAAATAGA GGAGGAAAGA ATCGATTCGA TAAAAGATGA AATATTAGAA ATTAAAATGG
Caecomyces_sp._Brit4        TGAAGATAGT CTTGAGAAAG GGGAAATTGA AAAAGATAGA ATCGATTCAA TAAAAGACGA AGTGTTACAA ATTAAAGTAG
Anaeromyces_robustus        TGAAGATAGT ATTGAGAATG GGGAAATTGA AGACGAAAGA ATTAATTTTA TAAAAGACGA TATATTAGAA ATTAAATTAG
Anaeromyces_contortus       TGAAGATAGT ATTGAGAATG GGGAAATTGA AGAAGATAGA ATTGATATAA TAAAAGATGA TATATTGGAA ATCAAATTAG

                            161
Ruminococcus_albus          GTGTCATTAT TGCGAGATAT TCAAGATGCG ATCCTATAGA CGATATCAAG TCGGGATTTG AAGACATGAT CGATATGTTT
Pecoramyces_ruminantium     GAATAGTTAT TGCTAGATAT TCAAGAGGGG ATTCGATGAA TGAACTAAAA CAAGAATTTG AAGAAATGAT TGATAGATTT
Piromyces_sp._FS3c          GAATAGTTAT TGCTAGATAC TCAAGAGGGG ATTCGATGAA TGAACTAAAA CGGGAATTTG AAGAAATGAT TGATAAATTT
Caecomyces_sp._Brit4        GAATAATTAT AGCTAAATAT TCAAGAGGTG ATCCTATTGA TACCATAAAG AAAGAATTTG AAGATATGAT TGATTCGTTT
Anaeromyces_robustus        GAATAATTAT AGCTAAATAT TCCAGAGGAG ATCCTATTTG TACAATAAAA AAAGAATTTG AAGATATGAT TGATTTTTTT
Anaeromyces_contortus       GAATAATTAT AGCTAAATAT TCAAGAGGTG ATCCTATTAA TATAATAAAG AAAGAGTTTG AGGATATGAT TGATTTGTTT

                            241
Ruminococcus_albus          TGCGAATCAT GGGACGGTGG TATATATGAA GATAATCTAT GGTTTGCATC ATTGGCTTAT CTTCTGGGGC TTGACAGCGC
Pecoramyces_ruminantium     TGTGAATCAT GGGACGGTGA AATCTATGAG GATAATCTTT GTTTTGCCTC TTTGGCCTAT CTTTTAGGAT TAGATGATGA
Piromyces_sp._FS3c          TGTGATTCAT GGGACGGTGA AATCTATGAG GATAATCTTT GTTTTGCCTC TTTGGCCTAT CTTTTAGGAT TAGATGATGA
Caecomyces_sp._Brit4        TGTGAATCTT GGGATGGTGA AATATATGAA GATAATCTTT GTTTTGCTTC CTTGGCATAT CTTTTAGGAC TAGATAATAC
Anaeromyces_robustus        TGTGAATTTT GGAATGGTGA AATATATGAA GATAATCTTT GGTTTGCTTC TTTGGCCTAT CTTTTAAAAT TAGATGAAGC
Anaeromyces_contortus       TGTGAATATT GGATTGGTGA AATATATGAA GATAATCTTT GTTTTGCTTC TTTAGCATAC CTTTTAGCAT TAGATGATAC

                            321
Ruminococcus_albus          AAAACTTGAA AAAATAAGGA AAAAATTGAT GGAAAGCGAC ACGTATGACT ATCTTATTGA TTTTATCCTA TCGGGTACTG
Pecoramyces_ruminantium     GCAATTGAAT AGAATAAGAA ATAAATTAAA AGAATCGGAT ACCTATGATT TTCTTATAGA TTTTGTACTT GTGGGTTTTG
Piromyces_sp._FS3c          GCAATTGAAT AGAATAAGAA ATAAATTAAA AGAATCGGAT ACCTATGATT TTCTTATAGA TTTTGTACTT GTGGGTTTTG
Caecomyces_sp._Brit4        ACAGCTAAAT AGGATAAGAA ATAAATTAAA GGAATCTGAT ACATACGATT ATCTTATAGA TTTTGTACTT GTAGGCGTTG
Anaeromyces_robustus        ATTGTTAAAA AGAATAAGAA ATAAATTAAA AGAATCTGAT ACATATGATT ATCTTATAGA TTTTATACTT ATAGGTTTTG
Anaeromyces_contortus       CTTATTAAAT AAAATAAGAA ATAAATTAAA AGAATCTGAT ACATATGATT ATCTGATAGA TTTTATATTG GTAGGTTTTG

                            401
Ruminococcus_albus          AAAGCAAGTT TGACAACAGT AAGATATCTT TCCCGCGTTC ATATAAAAAA CTGGTCAAAA GCATAAATGA GAATGACAAA
Pecoramyces_ruminantium     ATGATACTCT GGATGAATCA AAAATATCGT TTCCACGTCC CTATAAACAA TTGTTAAAGA GTATTAATGG TAAGGATAGA
Piromyces_sp._FS3c          ATGATACTCT GGATGAATCA AAAATATCGT TTCCACGTCC CTATAAACAA TTGTTAAGGA GTATTAATGG TAAGGATAGA
Caecomyces_sp._Brit4        ATGATGCTCA GGATAATTCA GAAATATCAT TCCCTCGTCC TTATAAACAA TTATTAAAAT GTATTAATGG TGAAGACAAA
Anaeromyces_robustus        ATGATTCTCA AGATAATTTA AAAATATCAT TTCCTCGTCC TTATAAAAAA TTATTAAAAT GTATTAATGG TCAAGATAGG
Anaeromyces_contortus       ATGATGATCA AAATGATTCA AAAATATCAT TTCCTCGTCC TTATAAAAAA TTGTTAAAAA GTATAAATGG TAAAGATAGA

                            481
Ruminococcus_albus          GAGTCGTTAT TAAAATATCT GCGCGGCTGG TATAAAGGCA GTCAGGAAAG TTCCTGGTAT GATACGCACA AGATCACAGA
Pecoramyces_ruminantium     GTTGCTTTCC AGAAATATCT ACGAGGTTGG TATAAAGGTT GTCAGGAAAG TTCGTGGTAT GATACTCATA AAATTGAAGA
Piromyces_sp._FS3c          GTTGCTTTCC AGAAATATCT ACGAGGTTGG TATAAAGGTT GTCAGGAAAG CTCGTGGTAT GATACTCATA AAATTGAAGA
Caecomyces_sp._Brit4        TATGCTTTTA AAAAATATTT ACGAGGCTGG TATAAAGGTT GTCAGGAAAG TTCATGGTAT GATACACACA AAATTGAAGA
Anaeromyces_robustus        GAAGCATTAT TAAAATACTT ACGAGGCTGG TATAAAGGTT GTCAAGAAAG CTCATGGTAT GATACACACA AAATTAAAGA
Anaeromyces_contortus       AATGCTTTTT TGAAGTATTT ACGAGGTTGG TATAAAGGTT GTCAGGAAAG CTCATGGTAC GATACACACA AAATTAAAGA

                            561
Ruminococcus_albus          CGACAACCTT TATTACGGCT ACTGGTGCTT TGATGCAGGT GCAGTTGCAA AAAGGCTTGG TCTTGAAGAC AGTGACCTGC
Pecoramyces_ruminantium     TGATAATCTA TATTTTGGAT ATTGGTGCTT TGAAGCCGGT GCTGTAGCTA AAAGACTAGG TTTTAAAGAC GATGATTTAA
Piromyces_sp._FS3c          TGACAATCTA TATTTTGGAT ATTGGTGCTT TGAAGCCGGT GCTGTAGCTA AAAGACTAGG TTTTAAAGAC GATGATTTAA
Caecomyces_sp._Brit4        TGATAATTTG TATTTTGGTT ATTGGTGTTT CGAAGCAGGA GCTGTAGCCA AAAGACTTGG TTTTAAAGAT GATGATTTAC
Anaeromyces_robustus        TGATAATTTA TATTTCGGAT ATTGGTGTTT TGAAGCAGGT GCTGTAGCTA AAAGACTTGG TTTTGAAGAC GATGATTTAA
Anaeromyces_contortus       TGACAATATA TATTTTGGGT ATTGGTGCTT TGAAGCAGGT GCTATAGCTA AAAGACTTGG TTTTAAAGAC GATGATTTAA

                            641
Ruminococcus_albus          AAAATGAACA GTACTATCCT TATGATATCG TTCATTTCAG ---CTGA
Pecoramyces_ruminantium     AAAATGAGCA ATATTATCCA TATGATATGG TTCATTTTAC ---ATGA
Piromyces_sp._FS3c          AAAATGAGCA TTATTATCCA TATGATATGG TTCATTTTGC ---ATGA
Caecomyces_sp._Brit4        AAAATGAGCA GTATTATCCA TATGATATGG TTCATTTTAC TGAATGA
Anaeromyces_robustus        AAAATAAACA ATATTATCCA TATGATATGA TTCATTATAT TGAATAA
Anaeromyces_contortus       AAAATGAACA GTATTATCCA TACGATATGG TTCATTTTAC TGAATGA
```

#### Text S2

**Expected PoNi amino acid sequence in *Anaeromyces robustus***

MRDTLRTKEYFDTFILEELEDIKMFEDSIENGEIEDERINFIKDDILEIKLGIIIAKYSRGDPICTIKKEFEDMIDF
FCEFWNGEIYEDNLWFASLAYLLKLDEALLKRIRNKLKESDTYDYLIDFILIGFDDSQDNLKISFPRPYKKLLKCIN
GQDREALLKYLRGWYKGCQESSWYDTHKIKDDNLYFGYWCFEAGAVAKRLGFEDDDLKNKQYYPYDMIHYIE

**Expected PoNi amino acid sequence in *Anaeromyces contortus***

MKTRDTLRSKEYFDTFILEELEDIKMYEDSIENGEIEEDRIDIIKDDILEIKLGIIIAKYSRGDPINIIKKEFEDMI
DLFCEYWIGEIYEDNLCFASLAYLLALDDTLLNKIRNKLKESDTYDYLIDFILVGFDDDQNDSKISFPRPYKKLLKS
INGKDRNAFLKYLRGWYKGCQESSWYDTHKIKDDNIYFGYWCFEAGAIAKRLGFKDDDLKNEQYYPYDMVHFTE

**Expected PoNi amino acid sequence in *Pecoramyces ruminantium***

MKMRDTLRTKEYFDTFIQEELEDIHMFEDSLSKGEIEEERIDSIKDEILEIKMGIVIARYSRGDSMNELKQEFEEMI
DRFCESWDGEIYEDNLCFASLAYLLGLDDEQLNRIRNKLKESDTYDFLIDFVLVGFDDTLDESKISFPRPYKQLLKS
INGKDRVAFQKYLRGWYKGCQESSWYDTHKIEDDNLYFGYWCFEAGAVAKRLGFKDDDLKNEQYYPYDMVHFT

**Expected PoNi amino acid sequence in *Caecomyces sp.* Brit4**

MKMRDTLRTKEYFDKFILEELEDIQMFEDSLEKGEIEKDRIDSIKDEVLQIKVGIIIAKYSRGDPIDTIKKEFEDMI
DSFCESWDGEIYEDNLCFASLAYLLGLDNTQLNRIRNKLKESDTYDYLIDFVLVGVDDAQDNSEISFPRPYKQLLKC
INGEDKYAFKKYLRGWYKGCQESSWYDTHKIEDDNLYFGYWCFEAGAVAKRLGFKDDDLQNEQYYPYDMVHFTE

**Expected PoNi amino acid sequence in *Piromyces sp.* FS3c**

MRDTLRTKEYFDTFIQEELEDIHMFEDSLSKGEIEEERIDSIKDEILEIKMGIVIARYSRGDSMNELKREFEEMIDK
FCDSWDGEIYEDNLCFASLAYLLGLDDEQLNRIRNKLKESDTYDFLIDFVLVGFDDTLDESKISFPRPYKQLLRSIN
GKDRVAFQKYLRGWYKGCQESSWYDTHKIEDDNLYFGYWCFEAGAVAKRLGFKDDDLKNEHYYPYDMVHFA

**Expected PoNi amino acid sequence in *Ruminococcus albus***

GenBank accession (nucleotide): ADKM02000110.1; GenBank accession (protein): EGC02125.1

MQMRDKLRKKEYFDTFIEEEIEDIQMFEDSLADGEIEEDRIDSIKDEILLIKLGVIIARYSRCDPIDDIKSGFEDMI
DMFCESWDGGIYEDNLWFASLAYLLGLDSAKLEKIRKKLMESDTYDYLIDFILSGTESKFDNSKISFPRSYKKLVKS
INENDKESLLKYLRGWYKGSQESSWYDTHKITDDNLYYGYWCFDAGAVAKRLGLEDSDLQNEQYYPYDIVHFS

**PoNi amino acid sequence alignment**

```
                            1
Ruminococcus_albus          MQMRDKLRKK EYFDTFIEEE IEDIQMFEDS LADGEIEEDR
Caecomyces_sp._Brit4        MKMRDTLRTK EYFDKFILEE LEDIQMFEDS LEKGEIEKDR
Pecoramyces_ruminantium     MKMRDTLRTK EYFDTFIQEE LEDIHMFEDS LSKGEIEEER
Piromyces_sp._FS3c          --MRDTLRTK EYFDTFIQEE LEDIHMFEDS LSKGEIEEER
Anaeromyces_robustus        --MRDTLRTK EYFDTFILEE LEDIKMFEDS IENGEIEDER
Anaeromyces_contortus       MKTRDTLRSK EYFDTFILEE LEDIKMYEDS IENGEIEEDR

                           41
Ruminococcus_albus          IDSIKDEILL IKLGVIIARY SRCDPIDDIK SGFEDMIDMF
Caecomyces_sp._Brit4        IDSIKDEVLQ IKVGIIIAKY SRGDPIDTIK KEFEDMIDSF
Pecoramyces_ruminantium     IDSIKDEILE IKMGIVIARY SRGDSMNELK QEFEEMIDRF
Piromyces_sp._FS3c          IDSIKDEILE IKMGIVIARY SRGDSMNELK REFEEMIDKF
Anaeromyces_robustus        INFIKDDILE IKLGIIIAKY SRGDPICTIK KEFEDMIDFF
Anaeromyces_contortus       IDIIKDDILE IKLGIIIAKY SRGDPINIIK KEFEDMIDLF

                           81
Ruminococcus_albus          CESWDGGIYE DNLWFASLAY LLGLDSAKLE KIRKKLMESD
Caecomyces_sp._Brit4        CESWDGEIYE DNLCFASLAY LLGLDNTQLN RIRNKLKESD
Pecoramyces_ruminantium     CESWDGEIYE DNLCFASLAY LLGLDDEQLN RIRNKLKESD
Piromyces_sp._FS3c          CDSWDGEIYE DNLCFASLAY LLGLDDEQLN RIRNKLKESD
Anaeromyces_robustus        CEFWNGEIYE DNLWFASLAY LLKLDEALLK RIRNKLKESD
Anaeromyces_contortus       CEYWIGEIYE DNLCFASLAY LLALDDTLLN KIRNKLKESD

                          121
Ruminococcus_albus          TYDYLIDFIL SGTESKFDNS KISFPRSYKK LVKSINENDK
Caecomyces_sp._Brit4        TYDYLIDFVL VGVDDAQDNS EISFPRPYKQ LLKCINGEDK
Pecoramyces_ruminantium     TYDFLIDFVL VGFDDTLDES KISFPRPYKQ LLKSINGKDR
Piromyces_sp._FS3c          TYDFLIDFVL VGFDDTLDES KISFPRPYKQ LLRSINGKDR
Anaeromyces_robustus        TYDYLIDFIL IGFDDSQDNL KISFPRPYKK LLKCINGQDR
Anaeromyces_contortus       TYDYLIDFIL VGFDDDQNDS KISFPRPYKK LLKSINGKDR

                          161
Ruminococcus_albus          ESLLKYLRGW YKGSQESSWY DTHKITDDNL YYGYWCFDAG
Caecomyces_sp._Brit4        YAFKKYLRGW YKGCQESSWY DTHKIEDDNL YFGYWCFEAG
Pecoramyces_ruminantium     VAFQKYLRGW YKGCQESSWY DTHKIEDDNL YFGYWCFEAG
Piromyces_sp._FS3c          VAFQKYLRGW YKGCQESSWY DTHKIEDDNL YFGYWCFEAG
Anaeromyces_robustus        EALLKYLRGW YKGCQESSWY DTHKIKDDNL YFGYWCFEAG
Anaeromyces_contortus       NAFLKYLRGW YKGCQESSWY DTHKIKDDNI YFGYWCFEAG

                          201
Ruminococcus_albus          AVAKRLGLED SDLQNEQYYP YDIVHFS-
Caecomyces_sp._Brit4        AVAKRLGFKD DDLQNEQYYP YDMVHFTE
Pecoramyces_ruminantium     AVAKRLGFKD DDLKNEQYYP YDMVHFT-
Piromyces_sp._FS3c          AVAKRLGFKD DDLKNEHYYP YDMVHFA-
Anaeromyces_robustus        AVAKRLGFED DDLKNKQYYP YDMIHYIE
Anaeromyces_contortus       AIAKRLGFKD DDLKNEQYYP YDMVHFTE
```

### Supplementary Tables

**Table S1. Nucleic acid sequence identities between the PoNi coding sequences, presented as a percent identity matrix.**

| | *A. robustus* | *A. contortus* | *O. sp.* strain C1A | *P. ruminantium* | *C. sp.* Brit4 | *P. sp.* FS3c | *R. albus* |
|---|---|---|---|---|---|---|---|
| *A. robustus* | 100.00% | 88.36% | 84.06% | 84.06% | 86.32% | 83.70% | 72.08% |
| *A. contortus* | 88.36% | 100.00% | 84.28% | 84.28% | 87.34% | 82.97% | 72.49% |
| *Pecoramyces ruminantium* | 84.06% | 84.28% | 100.00% | 100.00% | 84.86% | 97.66% | 73.39% |
| *C. sp*. Brit4 | 86.32% | 87.34% | 84.86% | 84.86% | 100.00% | 82.82% | 74.09% |
| *P. sp*. FS3c | 83.70% | 82.97% | 97.66% | 97.66% | 82.82% | 100.00% | 72.81% |
| *R. albus* | 72.08% | 72.49% | 73.39% | 73.39% | 74.09% | 72.81% | 100.00% |

**Table S2. Amino acid sequence identities between the PoNi proteins, presented as a percent identity matrix.**

| | *A. robustus* | *A. contortus* | *O. sp.* strain C1A | *P. ruminantium* | *C. sp.* Brit4 | *P. sp.* FS3c | *R. albus* |
|---|---|---|---|---|---|---|---|
| *A. robustus* | 100.00% | 84.65% | 78.41% | 78.41% | 81.14% | 77.43% | 73.57% |
| *A. contortus* | 84.65% | 100.00% | 82.46% | 82.46% | 83.33% | 80.26% | 72.81% |
| *Pecoramyces ruminantium* | 78.41% | 82.46% | 100.00% | 100.00% | 85.09% | 96.48% | 73.13% |
| *C. sp*. Brit4 | 81.14% | 83.33% | 85.09% | 85.09% | 100.00% | 82.46% | 75.44% |
| *P. sp*. FS3c | 77.43% | 80.26% | 96.48% | 96.48% | 82.46% | 100.00% | 71.37% |
| *R. albus* | 73.57% | 72.81% | 73.13% | 73.13% | 75.44% | 71.37% | 100.00% |

**Table S3. Brief summary of the top hit from BLAST searches, performed using blastp with default parameters against the non-redundant protein sequences (nr).**

| Query (Supplementary Text S2) | Top non-AGF hit accession | Coverage | E-value | Identity |
|---|---|---|---|---|
| *Anaeromyces robustus* | EGC02125.1 (*Ruminococcus albus*) | 99% | 1e-115 | 74.55% |
| *Anaeromyces contortus* | EGC02125.1 (*Ruminococcus albus*) | 99% | 8e-115 | 73.13% |
| *Pecoramyces ruminantium* | EGC02125.1 (*Ruminococcus albus*) | 100% | 1e-118 | 73.13% |
| *Caecomyces sp.* Brit4 | EGC02125.1 (*Ruminococcus albus*) | 99% | 7e-121 | 75.77% |
| *Piromyces sp*. FS3c | EGC02125.1 (*Ruminococcus albus*) | 100% | 1e-115 | 72.00% |

#### Supplementary Figure

**Figure S1.** Full, expanded maximum-likelihood phylogenetic tree of fungal and bacterial PoNi proteins. The fungal proteins are shown in bold. This phylogenetic tree only includes five fungal PoNi proteins, as we could not find PoNi homologs in other fungal lineages. Shimodaira-Hasegawa approximate-likelihood ratio test (aLRT SH-like) values are provided as estimates of branch support next to the nodes. GenBank accession numbers are shown in brackets. The tree is drawn to scale, with branch lengths measured in the number of substitutions per site.

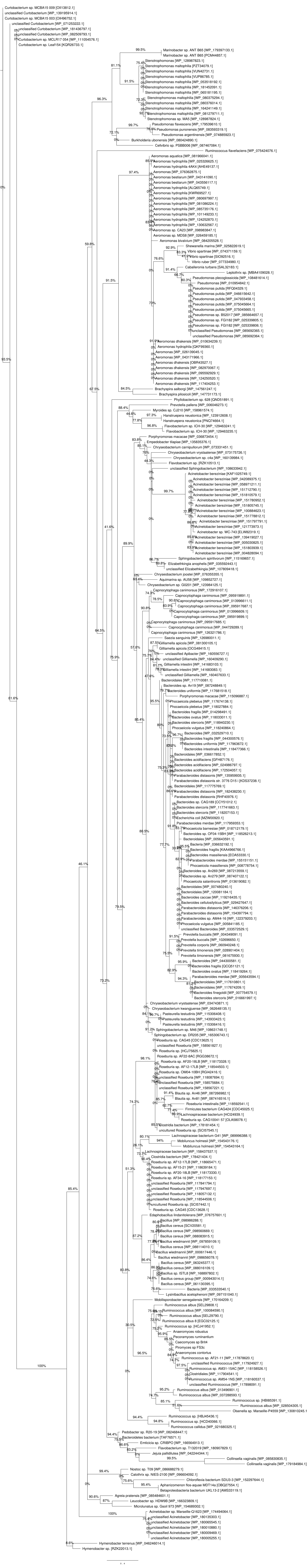
